# Human Topoisomerase IIα Promotes Chromatin Condensation Via a Phase Transition

**DOI:** 10.1101/2024.10.15.618281

**Authors:** Meiling Wu, Curtis Beck, Joyce H. Lee, Robert M. Fulbright, Joshua Jeong, James T. Inman, Mitchell V. Woodhouse, James M. Berger, Michelle D. Wang

## Abstract

Topoisomerase II (topo II) enzymes are essential enzymes known to resolve topological entanglements during DNA processing. Curiously, while yeast expresses a single topo II, humans express two topo II isozymes, topo IIα and topo IIβ, which share a similar catalytic domain but differ in their intrinsically disordered C-terminal domains (CTDs). During mitosis, topo IIα and condensin I constitute the most abundant chromosome scaffolding proteins essential for chromosome condensation. However, how topo IIα enables this function is poorly understood. Here, we discovered a new and functionally distinct role for human topo IIα – it condenses DNA and chromatin at a low topo IIα concentration (100 pM or less) during a polymer-collapse phase transition. The removal of the topo IIα CTDs effectively abolishes its condensation ability, indicating that the condensation is mediated by the CTDs. Although topo IIβ can also perform condensation, it is about 4-fold less effective. During the condensation, topo IIα-DNA condensates form along DNA, working against a DNA tension of up to 1.5 pN, greater than that previously reported for yeast condensin. In addition, this condensation does not require ATP and thus is independent of topo IIα’s catalytic activity. We also found that condensation and catalysis can concurrently proceed with minimal mutual interference. Our findings suggest topo IIα may directly participate in chromosome condensation during mitosis.

---

Fundamental processes that occur over DNA, such as transcription and replication, generate torsional stress and change DNA topology due to the helical nature of DNA^1-7^. DNA supercoiling, catenation, and topological entanglements arising from these processes are resolved by topoisomerases, ubiquitous enzymes found in all branches of life^8-12^. Type IIA topoisomerases (topo IIs) are essential enzymes for all organisms and function by an ATP-dependent strand passage mechanism, whereby topo II creates a transient double-stranded break in one DNA segment, passes a second DNA segment through this break, and subsequently religates the DNA break^13-15^.

Curiously, vertebrates, such as humans, express two topo II isoforms, topo IIα and topo IIβ, while invertebrates, like yeast, express a single isoform. Human topo IIα and topo IIβ share significant sequence and structural homology in their core domains for DNA binding and catalytic activity, but they have distinct sequences in their C-terminal domains (CTDs)^16-18^. Both topo IIα and topo IIβ are essential for vertebrate development. Deletion of topo IIα results in lethality during embryonic development; deletion of topo IIβ allows development in utero but results in perinatal death^8,19-21^. In contrast, human cell lines show differential requirements for the two topo II isoforms. Deletion of human topo IIα, but not topo IIβ, causes mitotic failure and results in cell death, suggesting an essential role for topo IIα in mitotic functions^9,16,22^. Supporting this notion, human topo IIα expression and degradation are highly cell cycle regulated, with its presence restricted to the late-S to M phase^11,23,24^. In contrast, topo IIβ is present at uniform levels throughout the cell cycle^11,24^. During mitosis, human topo IIα and condensin I constitute the most abundant and crucial chromosome scaffolding proteins required for mitotic chromosome condensation, and topo IIα performs the critical role of decatenating the chromosomes^8,25-30^.

Thus far, cellular and biochemical studies have suggested that the functional differences between topo IIα and topo IIβ are dictated, at least in part, by the CTDs of the two topo IIs^9,16,31-37^(Supplementary Fig. 1). The topo II CTD is notable in that it consists of a large, typically 300-400 amino acid intrinsically disordered element that has high sequence complexity^17,38^. A recent study has found that yeast topo II and both human topo IIα and topo IIβ can phase separate to form condensed bodies and that the presence of DNA stimulates this separation and further increases the fluidity of the condensates to form liquid droplets^17^.

Although human topo IIα, but not topo IIβ, is essential for mitosis, it is unclear whether topo IIα and topo IIβ have fundamentally different innate actions on chromatin at a molecular level that account for their differential roles. It has been reported that topo IIα and topo IIβ differ in their abilities to relax positively *versus* negatively supercoiled naked DNA^39-41^. However, it is unknown whether topo IIα is more effective than topo IIβ at decatenating chromatin, which would be important for assisting chromosome segregation. Here, we investigate the functional differences between topo IIα and topo IIβ, which could provide insights into their differential roles in mitosis.

## Topo IIα is somewhat more efficient than topo IIβ at decatenating chromatin

DNA replication completion requires decatenation of the intertwined chromosomes behind the replication fork to allow chromosome segregation. In humans, topo IIα expression increases towards the late S-phase^24^, suggesting a role of topo IIα in facilitating chromosome decatenation to ensure the segregation of the two daughter chromosomes. However, experiments examining how topo IIα acts on a catenated chromatin substrate have been lacking. To determine the efficiency of topo IIα in relaxing an intertwined chromatin substrate, we employed a magnetic tweezers-based method we previously developed to assess the efficiency by which topo II relaxes a chromatinized DNA substrate^42^. In this experiment (Fig. 1a), a chromatin substrate was first torsionally constrained between a magnetic bead and the surface of a microscope coverslip and held under 0.5 pN force. Next, human topo IIα or topo IIβ was added, and the chromatin tether was supercoiled by rotating the magnetic bead at 3.6 turns/s, a value close to the replication rate reported in human cells^43-45^. We then used the DNA extension to assess the ability of topo II to relax the supercoiling during an introduction of 1000 turns. We expected that if topo II could fully keep up with the rotation during this time, the DNA extension would remain near its initial length; otherwise, it would reduce with continued rotation.

**Figure 1.**
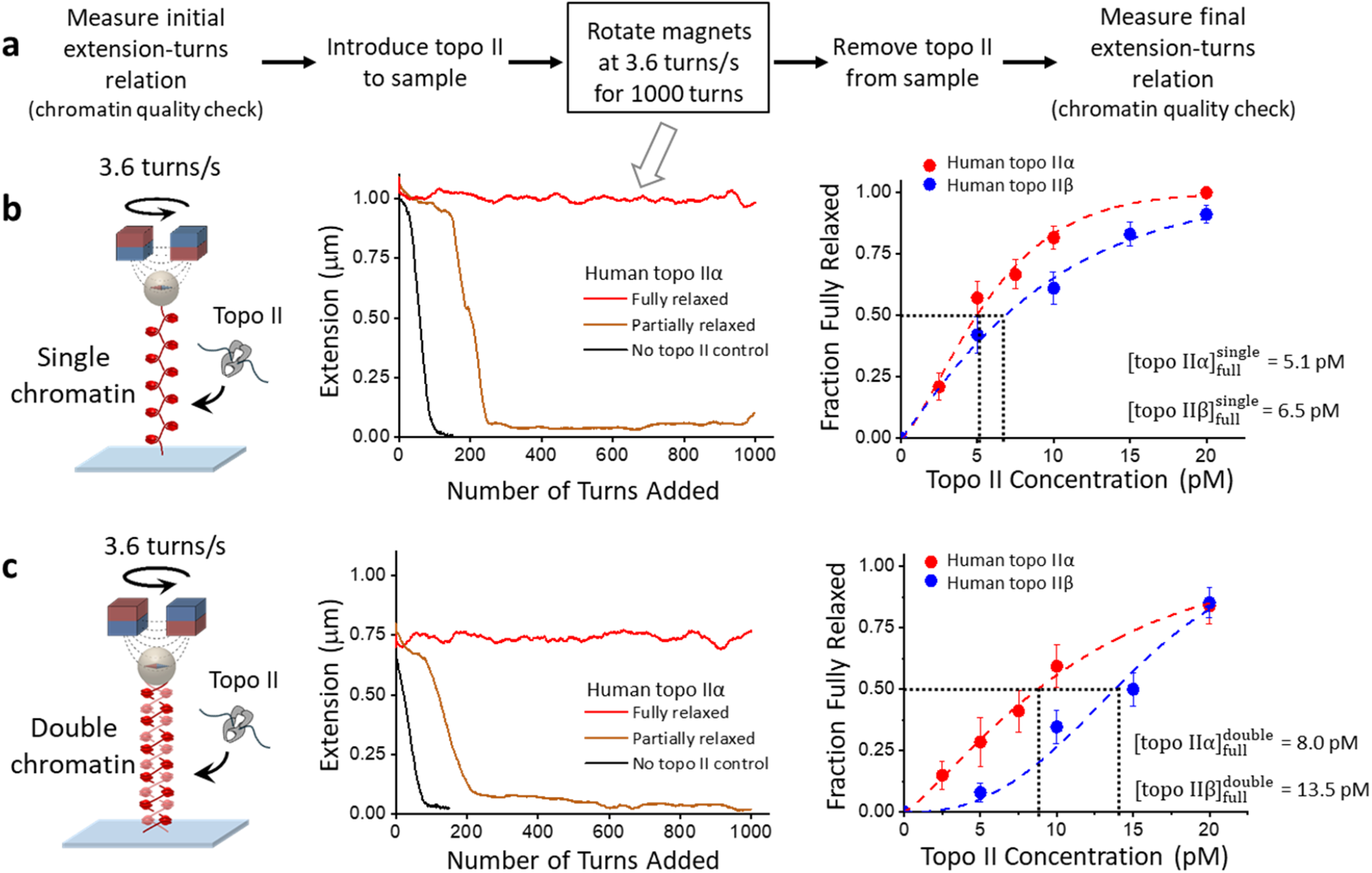
Human topo IIα is 1.7X more efficient than topo IIβ in chromatin decatenation. Topo II catalytic activity with chromatin fibers was assessed with a continuous winding assay using a magnetic tweezers (MT) setup. Each chromatin fiber contained ∼50 nucleosomes on average. The chromatin substrate was torsionally constrained between a magnetic bead and the surface of a microscope coverslip. **a**. Experimental scheme of the continuous winding assay. The initial and final extension-turns relations of the chromatin fibers were measured in the absence of topo II to quantify the saturation, geometry, and stability of the chromatin fibers. After introducing topo II, the chromatin substrate was supercoiled via the paramagnetic bead by rotating a pair of magnets at a constant rate of 3.6 turns/s to mimic the rate of supercoiling generation during replication in human cells^43-45^. **b**. Comparison of the efficiency of human topo IIα versus topo IIβ in relaxing single-chromatin fibers. The middle panel shows example traces from human topo IIα. A trace is identified to be a fully relaxed trace (red) when the topo II activity was able to remove > 95% of the supercoiling introduced by the constant magnet rotation, resulting in the extension remaining near the initial value. The right panel shows the fraction of fully relaxed traces as a function of topo II concentration. 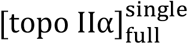 and 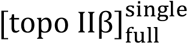 are the concentrations at which 50% of traces are fully relaxed. These values reflect the effectiveness of topo II in relaxing a single-chromatin substrate. Each data point is calculated from ∼50 traces; error bars indicate the counting errors. **c**. Comparison of the efficiency of human topo IIα versus topo IIβ in relaxing double-chromatin fibers. 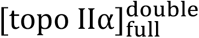 and 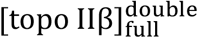 are the concentrations at which 50% of traces are fully relaxed. These values reflect the effectiveness of topo II in relaxing a double-chromatin substrate. Each data point is calculated from ∼40 traces; error bars indicate the counting errors.

Using this method, we compared the ability of topo IIα to fully relax a supercoiled single-chromatin substrate (which would form in front of the replisome) with its ability to fully relax an intertwined double-chromatin substrate (which would form behind the replisome). The chromatin fibers in the single and double-chromatin substrates have identical DNA lengths (Methods). We compared the activity of topo IIα with that of topo IIβ. Our data show that the fraction of fully relaxed tethers increased with an increase in topo II concentration, which was true for both single and double-chromatin substrates of either human topo IIα or topo IIβ (Fig. 1b,c). We characterized the effectiveness of topo II in relaxing a substrate using the topo II characteristic concentration [topo IIα]_full_, at which 50% of tethers of a given type of substrate was fully relaxed. The lower the value of [topo IIα]_full_, the more efficient the enzyme is in removing DNA supercoils or decatenation since less topo II is required to achieve full relaxation. We found that the characteristic concentration of topo IIα on the single-chromatin substrate was about 75% of that on the double-chromatin substrate 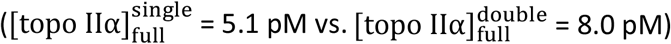. By comparison, the characteristic concentration topo IIβ on a single-chromatin substrate was nearly 2-fold less than on the double-chromatin substrate 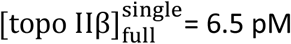 and 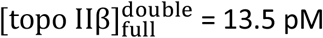. Overall, our results show that both human topo IIα and topo IIβ are more efficient at relaxing a single-chromatin substrate than a double-chromatin substrate, a finding consistent with previous data obtained for yeast topo II^42^. Our previous chromatin torsional mechanical studies suggest that DNA supercoiling generated during replication elongation is preferentially directed to the front of the replication fork, where there is a single-chromatin substrate during replication elongation^42^. Thus, efficient relaxation of the single-chromatin substrate by human topo IIα (and IIβ) ensures that the chromatin torsional mechanical properties and topoisomerase activities work synergistically during replication elongation (and transcription). During the late S-phase, as replication approaches termination, the single-chromatin substrate in front of the replisome diminishes, leaving behind an intertwined double-chromatin substrate, which is less efficient for topo II to relax. We also found that human topo IIα is 1.7X more efficient than human topo IIβ at decatenating an intertwined double-chromatin substrate, suggesting topo IIα is somewhat more effective at facilitating replication completion and subsequent chromosome segregation. Our data may provide a plausible explanation for the need to elevate the expression of human topo IIα during the late S-phase.

## Topo IIα unexpectedly drives efficient chromatin compaction

Our observation that human topo IIα is more effective than topo IIβ at decatenating braided chromatin suggests human topo IIα could facilitate replication completion. However, it does not explain the essential need for topo IIα in cell viability, given that human topo IIβ can also decatenate chromatin, albeit a little less effectively. This conundrum indicates that human topo IIα may perform a function distinct from its catalytic activity. Previously, topo IIα has been found to work with condensin I in scaffolding the chromosomes during mitotic chromosome condensation^8,25-29^. Although the exact nature of this role remains unclear, it has been suggested that condensin I drives the chromosome condensation by loop extrusion while topo IIα assists the topology of the condensation by catenating and decatenating chromosomes^17,27-29,33,46,47^.

Interestingly, in a previous study we found evidence that human topo IIα may be able to directly compact naked DNA^48^, suggesting that the enzyme may possess as-yet-undescribed physical properties outside of supercoil removal and DNA unlinking. At the time, we interpreted this compaction as resulting from an ability of topo IIα to bind to a DNA crossover and secure a small DNA loop in an ATP-independent manner. This interpretation did not require that human topo IIα molecules interact with each other but the data available at the time could not exclude this possibility. Assessing topo II-topo II interactions requires a real-time assay to monitor the compaction process rather than surveying the final compaction outcome, as was done in our previous study. To accomplish this goal, we used a magnetic tweezers-based assay similar to that used in Fig. 1 but now used DNA extension, which directly measures the extent of the DNA compaction, to monitor the entire process of DNA compaction starting from human topo IIα introduction (Fig. 2a). This is a high-throughput real-time assay where a couple hundred DNA tethers can be monitored simultaneously. Using this method, we systematically examined a broad range of topo II concentrations with a few example measurements shown in Fig. 2b-e.

**Figure 2.**
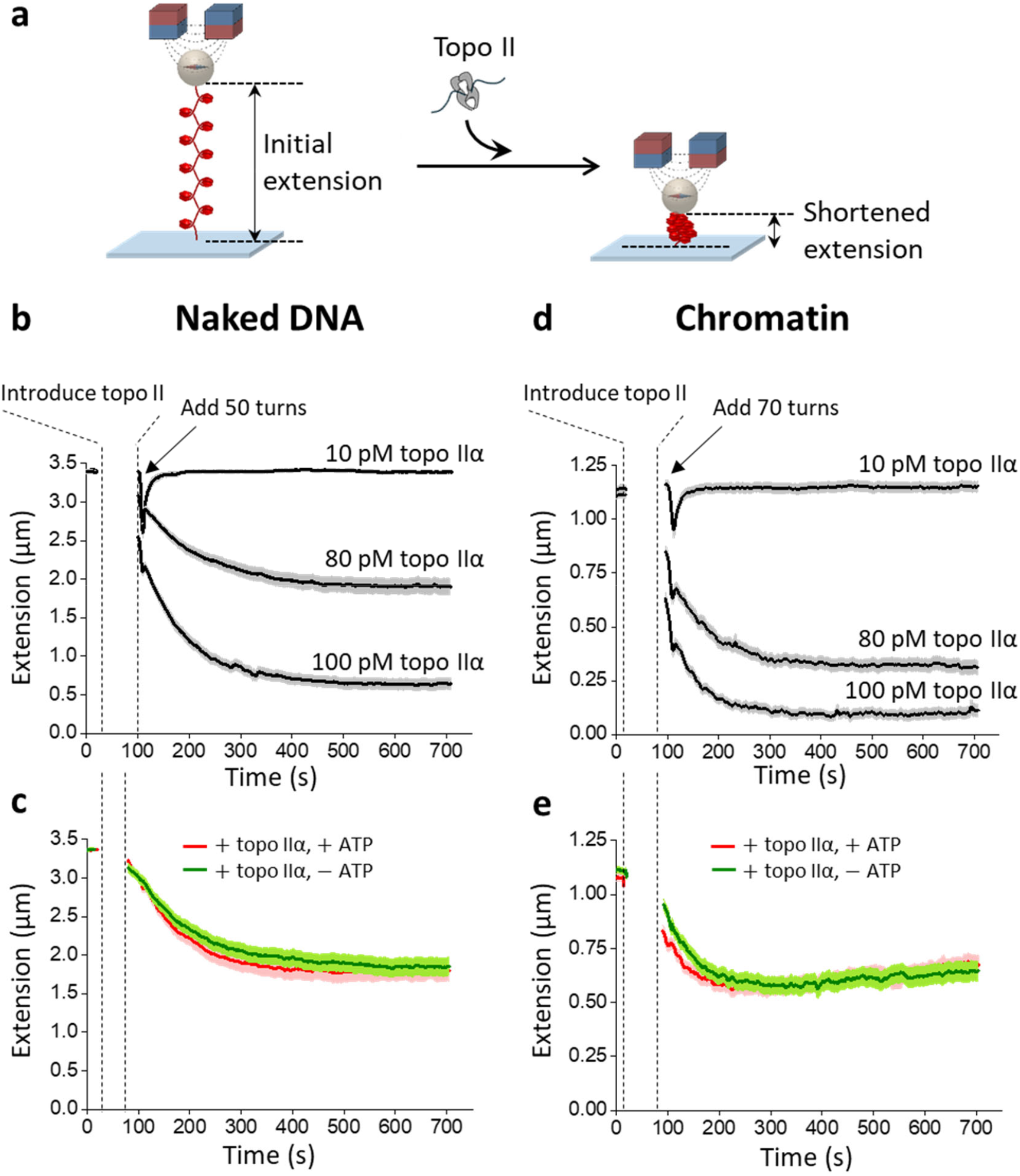
Direct measurement of chromatin compaction induced by human topo IIα. **a**. Experimental scheme for monitoring the chromatin compaction process. A chromatin substrate was torsionally constrained between a magnetic bead and the surface of a microscope coverslip. The extension of the chromatin tether, which directly measures the extent of the DNA compaction, was monitored before and after topo II introduction. **b**. Human topo IIα induces DNA compaction. DNA extension *versus* time in the presence of 10 pM, 80 pM, or 100 pM human topo IIα in topo reaction buffer with 1 mM ATP. Immediately after introducing topo II, 50 turns were added to the DNA to check topo IIα activity. Under all three topo concentrations, topo IIα removed all the turns rapidly. Subsequent tether shortening over time indicates topo II compaction of DNA. Each thick black curve is an average of ∼100 individual traces, with the shaded region indicating the SEM. **c**. DNA compaction induced by human topo IIα does not require ATP. DNA extension *versus* time in the presence of human topo IIα with and without ATP. These experiments were conducted under a condition where the final extension was at about 50% of the initial extension. Each curve represents an average of ∼60 traces, with the shaded region indicating the SEM. **d**. Human topo IIα induces chromatin compaction. Same as (b) except with a chromatin substrate. Each curve represents an average of ∼80 individual traces, with the shaded region indicating the SEM. **e**. Chromatin compaction by human topo IIα does not require ATP. Same as (c) except with a chromatin substrate. Each curve represents an average of ∼63 traces, with the shaded region indicating the SEM.

We first examined how human topo IIα compacts naked DNA, which has the same length as the DNA template used for nucleosome assembly but with a non-repetitive sequence replacing the 64 successive nucleosome positioning elements (Fig. 2b; Supplementary Fig. 2). After introducing topo IIα, we immediately performed a topo IIα activity check by rapidly adding 50 turns. In the absence of topo IIα, these extra turns were sufficient to buckle the DNA into a plectoneme^49-52^, shortening the DNA extension (Supplementary Fig. 3). In the presence of topo IIα, these extra turns only transiently perturbed the extension, and topo IIα removed these extra turns rapidly (Fig.2; Supplementary Fig. 4). Unexpectedly, the level of DNA compaction was seen to increase as the concentration of topo IIα was elevated. At a low topo IIα concentration (10 pM), the DNA remained at full extension over the observation time. By contrast, as the topo IIα concentration was increased, the extension decreased with time toward a final extension plateau, indicating DNA compaction. The higher the topo IIα concentration, the lower the final extension plateau, indicating more DNA compaction. The extension versus time plot could be well fit by a single decaying exponential with an offset (Supplementary Fig. 5). To investigate whether the compaction involves strand passage activity, we conducted experiments with and without ATP (Fig. 2c). To facilitate a direct comparison of the two conditions, we omitted the topo IIα activity check. Interestingly, we found no significant differences in the compaction kinetics with and without ATP, indicating that the compaction is driven by a process distinct from the catalytic activity^53^.

Although our experiments show that human topo IIα can compact naked DNA, it is unclear if it can also compact chromatin, a substrate more relevant for chromosome condensation. To investigate whether human topo IIα can compact chromatin, we performed corresponding experiments using a chromatin substrate. Since chromatin requires more turns to buckle than naked DNA, we added 70 turns instead of 50 for the initial topo IIα activity check (Fig. 2d) (importantly, this amount of torsion is insufficient to destabilize the nucleosomes; Supplementary Fig. 3)^42,54,55^. We found that human topo IIα also compacted chromatin, and the extent of the compaction again increased with an increase in topo IIα concentration. In addition, chromatin compaction also does not require ATP, since chromatin compacted similarly with and without ATP (Fig. 2e), further supporting the conclusion that compaction results from a process distinct from catalytic activity. Interestingly, we found that the compaction does require Mg^2+^ (Supplementary Fig. 6). In contrast, previous studies show that topo IIα-DNA liquid-liquid phase separation (LLPS) does not require Mg^2+^, suggesting that these two processes have distinct physical properties^17^.

## Topo IIα condenses chromatin via a phase transition

The observations in Fig. 2 suggest that topo IIα may condense the DNA via a polymer-collapse phase transition^56^, which occurs at a much lower topo IIα concentration than that required for LLPS. Such a phase transition is expected to be highly cooperative, with the DNA extension undergoing a sharp decrease to a condensed state once a critical topo II concentration [topo IIα]_c_ is reached, and the condensed state should have an extension close to zero. However, the value of [topo IIα]_c_ depends on temperature, force on the substrate, nucleosome density, buffer, salt, etc., which constitute system parameters of a complex phase diagram. So, even when measurements are conducted under specified target conditions, slight variations in these parameters can blur the measured transition. Nevertheless, the sharpness of a transition, together with the completeness of a condensed state, are excellent indicators of such a polymer-collapse phase transition.

To determine whether the compaction has features characteristic of such a polymer-collapse phase transition, we measured the final extension after compaction as a function of human topo IIα concentration for naked DNA (Fig. 3a) and chromatin (Fig. 3b). In each plot, there is a sharp reduction in the final extension when human topo IIα concentration reaches a critical value with the extension reaching the minimum attainable value for the instrument (Methods), indicating topo IIα condenses DNA or chromatin via a polymer-collapse phase transition. For simplicity, we fit each plot with an *n*-th order Hill equation, yielding a critical concentration [topo II]_c_ at which the DNA extension is reduced by half of the maximum extension reduction, and a Hill coefficient *n*. Since the lower the critical concentration, the more effective topo II is at inducing the phase transition, we use 1/[topo II]_c_ as a measure of the condensation efficiency (Fig. 3c). We found [topo IIα]_c_ = 78 pM and *n* = 10 for the naked DNA substrate (Fig. 3a) and 62 pM and *n* = 4 for the chromatin substrate (Fig. 3b). Thus, topo IIα can induce a phase transition in either DNA or chromatin at a low concentration, with the transition being highly cooperative.

**Figure 3.**
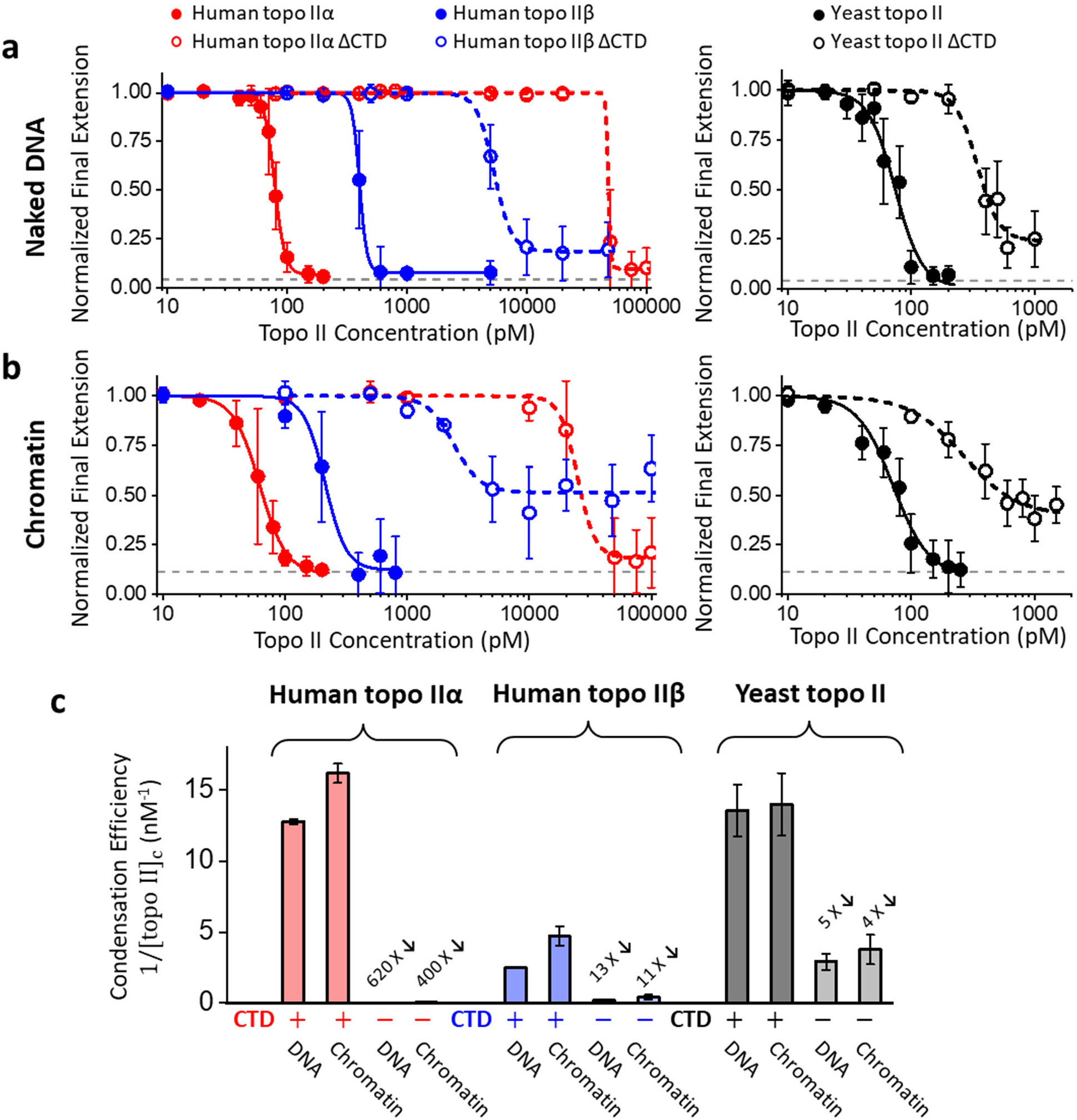
DNA condensation by topo IIs depends on their CTDs. **a**. Topo II condenses naked DNA by a phase transition. Each normalized final extension data point represents an average of over ∼100 individual traces, with error bars indicating SDs. The normalized final extension *versus* topo II concentration is fitted with a Hill equation: 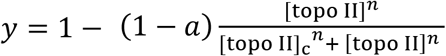, where *a* is the minimum normalized extension, [topo II]_c_ is the critical topo II concentration at which the extension is reduced by half, and *n* is the Hill coefficient. The dashed line indicates the minimum attainable extension by the instrument. **b**. Topo II condenses chromatin by a phase transition. Same as (a) except with chromatin substrates. Each data point represents an average of over ∼100 individual traces, with error bars indicating SDs. **c**. Condensation efficiency of all six topo IIs on naked DNA and chromatin. The condensation efficiency of topo II is characterized by 1/[topo II]_c_, which is obtained from Hill equation fitting. Error bars represent uncertainties within 95% confidence intervals. The reduction in the condensation efficiency of each CTD-truncated topo II relative to that of the full-length topo II on the same substrate is also indicated.

To directly compare the condensation abilities of human topo IIα and human topo IIβ, we repeated the experiments shown in Fig. 2 using human topo IIβ. Topo IIβ condensation also shows characteristics of a polymer-collapse phase transition. We found [topo IIβ]_c_ = 400 pM and *n* = 14 for the naked DNA substrate (Fig. 3a) and 213 pM and *n* = 5 for the chromatin substrate (Fig. 3b). Thus, human topo IIα is more effective in condensation than human topo IIβ: about 5 times more effective on the naked DNA substrate and about 3.5 times more effective on the chromatin substrate. We also found that topo IIα condenses naked DNA and chromatin with similar efficiencies, although chromatin condensation showed a less sharp transition, likely due to variations in the nucleosome density among tethers (Methods).

We further examined the condensation abilities of human topo IIα and topo IIβ that lacked their CTDs (Supplementary Fig. 1). Strikingly, we detected a greatly reduced condensation ability for topo IIα ΔCTD: [topo IIα ΔCTD]_c_ = 48,000 pM for the naked DNA substrate and 25,000 pM for the chromatin substrate (Fig. 3a,b,c). Thus, human topo IIα ΔCTD is about 600 X less effective at condensing DNA or chromatin; its ability to condense the substrate is essentially abolished. This observation shows that the CTD of human topo IIα is responsible for highly efficient condensation. Removing the CTD from the human topo IIβ also reduced its ability to condense DNA or chromatin by about 10-fold (Fig. 3a,b,c), suggesting that the CTD of human topo IIβ also contributes to condensation, albeit not as effectively as the CTD of human topo IIα. In addition, topo IIβ ΔCTD data, and to some extent the topo IIα ΔCTD data, show a significant non-zero plateau in the final extension of both the naked DNA and chromatin and a more gradual transition to the compacted state, indicating that the compaction process is less characteristic of a phase transition.

To explore the generality of our conclusions, we also examined yeast topo II using the experimental workflow shown in Fig. 2a. We found that yeast topo II can condense DNA and chromatin similarly to human topo IIα. However, the yeast topo II ΔCTD construct did not induce as dramatic a reduction in the condensation ability as the human topo IIα ΔCTD construct and its behavior is somewhat similar to that of topo IIβ. Thus, compared with the CTDs of human topo IIβ and yeast topo II, the CTD of human topo IIα is far more potent in contributing to the condensation ability.

Overall, human topo IIα and topo IIβ have markedly distinct abilities to condense DNA or chromatin after a polymer-collapse phase transition. Compared to human topo IIβ, human topo IIα is about 4 times more effective in condensing chromatin, owing nearly entirely to its CTD. This finding raises the possibility that human topo IIα can directly condense chromosomes during mitosis after a phase transition. If topo IIα condenses DNA after a phase transition, the critical topo IIα concentration should be sensitive to the force in the DNA. In our experiment, the DNA was held under a constant 0.5 pN force. A control experiment shows that the critical topo IIα concentration decreases with decreased force in the DNA (Supplementary Fig. 7), which is expected as the force in the DNA resists DNA condensation. *In vivo*, motor proteins, such as condensin, could, in principle, provide this force. The force used here is well within the range of the force generated by yeast condensin, which has been reported to be 0.4 – 1.0 pN^46^.

## Topo IIα condensation does not interfere with its catalytic activity

While topo IIα induces chromatin condensation, it is unclear if topo IIα can still continuously resolve supercoiling of a single-chromatin substrate or continuously decatenate an intertwined double-chromatin substrate, as would be required for mitosis. To investigate whether human topo IIα condensation of chromatin interferes with its ability to continuously relax a single-chromatin substrate (Fig. 4a) or double-chromatin substrate (Fig. 4b), we used the experimental configuration shown in Fig. 1, except using a higher human topo IIα concentration, which was near the critical concentration so that significant condensation could occur.

**Figure 4.**
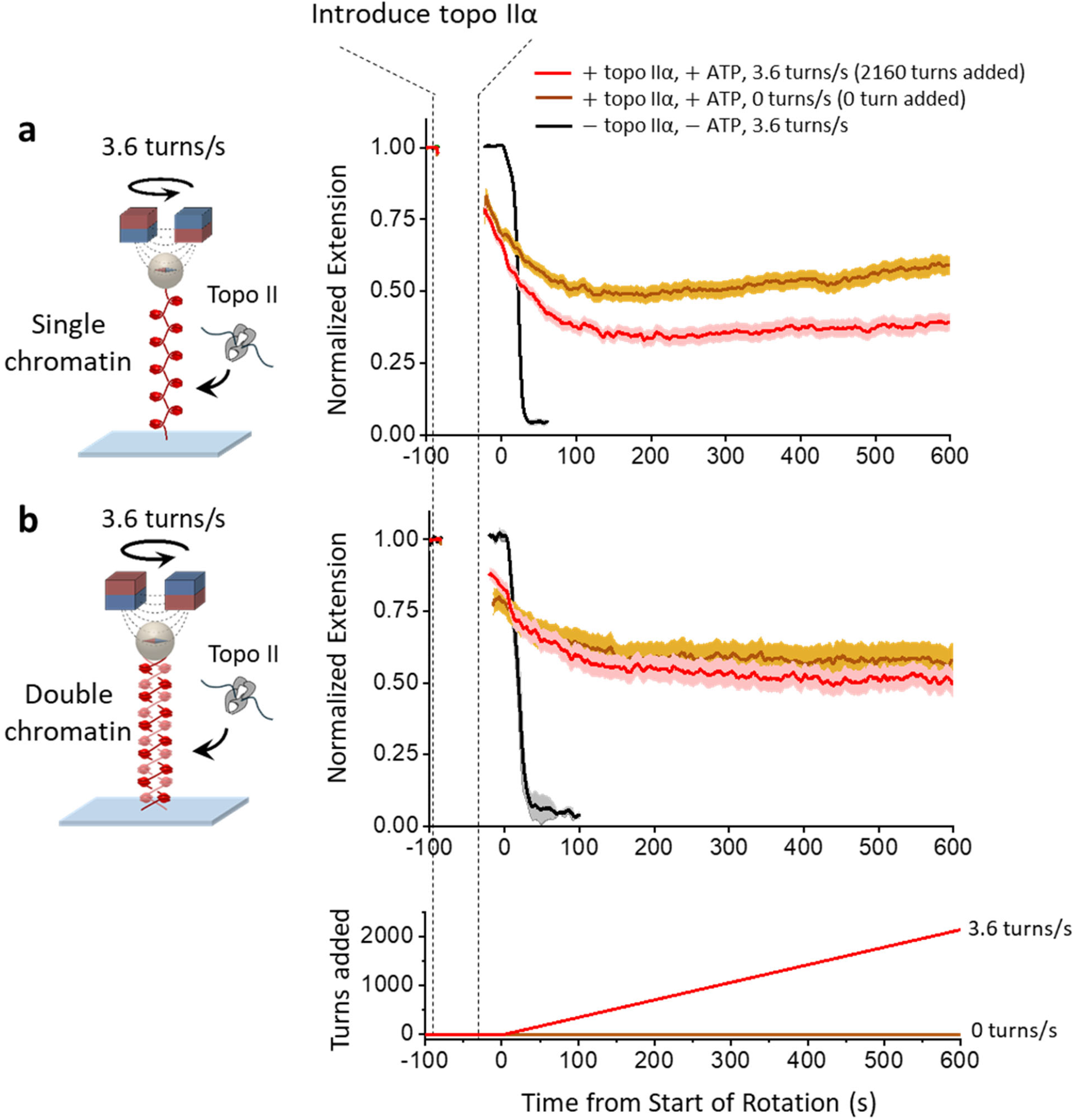
Human topo IIα chromatin condensation does not interfere with its catalytic activity. Concurrent topo IIα catalytic activity and topo IIα-induced chromatin condensation were assessed using the same experimental configuration as shown in Fig. 1b for single chromatin or Fig. 1c for double chromatin. Experiments were conducted under a condition where the final extension was at about 50% of the initial extension. **a**. Concurrent topo IIα supercoiling relaxation and condensation of the single-chromatin substrate. We measured the extension of a single-chromatin substrate as it was supercoiled at 3.6 turns/s to add 2160 turns (red). During this time, more than 98% of the added turns were removed by topo IIα (Methods). For comparison, the same experiment was conducted without adding any turns (brown). We also show a no-topo control (black). Each curve represents an average of ∼ 68 individual traces, with the shaded region indicating the SEM. **b**. Concurrent topo IIα chromatin decatenation and condensation of the double-chromatin substrate. Same as (a) except with the double-chromatin substrate. The bottom plot shows the turns added to the substrate. We measured the extension of a double-chromatin substrate as it was supercoiled at 3.6 turns/s to add 2160 turns (red). During this time, more than 99% of the added turns were removed by topo IIα (Methods). For comparison, the same experiment was conducted without adding any turns (brown). We also show a no-topo control (black). Each curve represents an average of ∼ 24 individual traces, with the shaded region indicating the SEM.

For each substrate, three experiments were conducted for comparison. During one control experiment in the absence of topo IIα, the extension was rapidly brought to the surface by continuous rotation of the magnets at 3.6 turns/s, because there was no relaxation of the introduced supercoiling. A second control experiment with topo IIα and ATP was conducted to assess the extent of chromatin condensation without supercoiling the substrate (no magnet rotation), so that the resulting extension shortening was solely due to topo IIα-induced chromatin condensation. The third and actual experiment was performed with topo IIα and ATP under continuous rotation of the magnets at 3.6 turns/s. The resulting extension shortening was due to a combined contribution from topo IIα-induced chromatin condensation and the residual supercoiling present in the substrate that was not relaxed by topo IIα. The extension plateaued at a lower value for the single-chromatin substrate than the double-chromatin substrate. This is likely because topo IIα substrate relaxation requires interaction with a DNA crossing, which more likely occurs on a single-chromatin substrate after the chromatin buckles to form a plectoneme, leading to a shortened extension^42,55^. In contrast, such a crossing occurs readily on a double chromatin substrate^42^. We estimate that topo IIα relaxed about 98% of 2160 turns introduced for the single-chromatin substrate and about 99% of 2160 turns introduced for the double-chromatin substrate (Methods). This demonstrates that topo IIα chromatin condensation minimally interferes with its ability to relax the chromatin substrates of a continuously supercoiled single-chromatin substrate or a continuously catenated double-chromatin substrate.

## Topo IIα generates a driving force to condense DNA

Since human topo IIα condenses DNA, it should generate a force to drive condensation. Although magnetic tweezers are ideally suited for experiments under a constant force, optical tweezers are more suited for stall force measurements. Thus, we performed experiments using a dual optical trap, tethering a lambda DNA molecule between the two traps (Fig. 5a). To begin the experiment, we extended the DNA to 6 μm, significantly smaller than its contour length (16.4 μm), and then held the trap positions fixed to allow topo IIα condensing DNA. As shown in Fig. 5b, at 2.5 nM topo II, the force in the DNA increased and eventually plateaued. We use the final force plateau as a measure of the condensation force: topo IIα generated about 0.8 pN, while topo IIβ generated about 0.4 pN, significantly weaker than that of topo IIα. As expected, topo IIα ΔCTD did not generate any detectable force. We also found that Mg^2+^, but not ATP, is required for the condensation force generation (Supplementary Fig. 8), again consistent with the requirement of Mg^2+^ for topo II*α*-induced chromatin condensation^57^.

**Figure 5.**
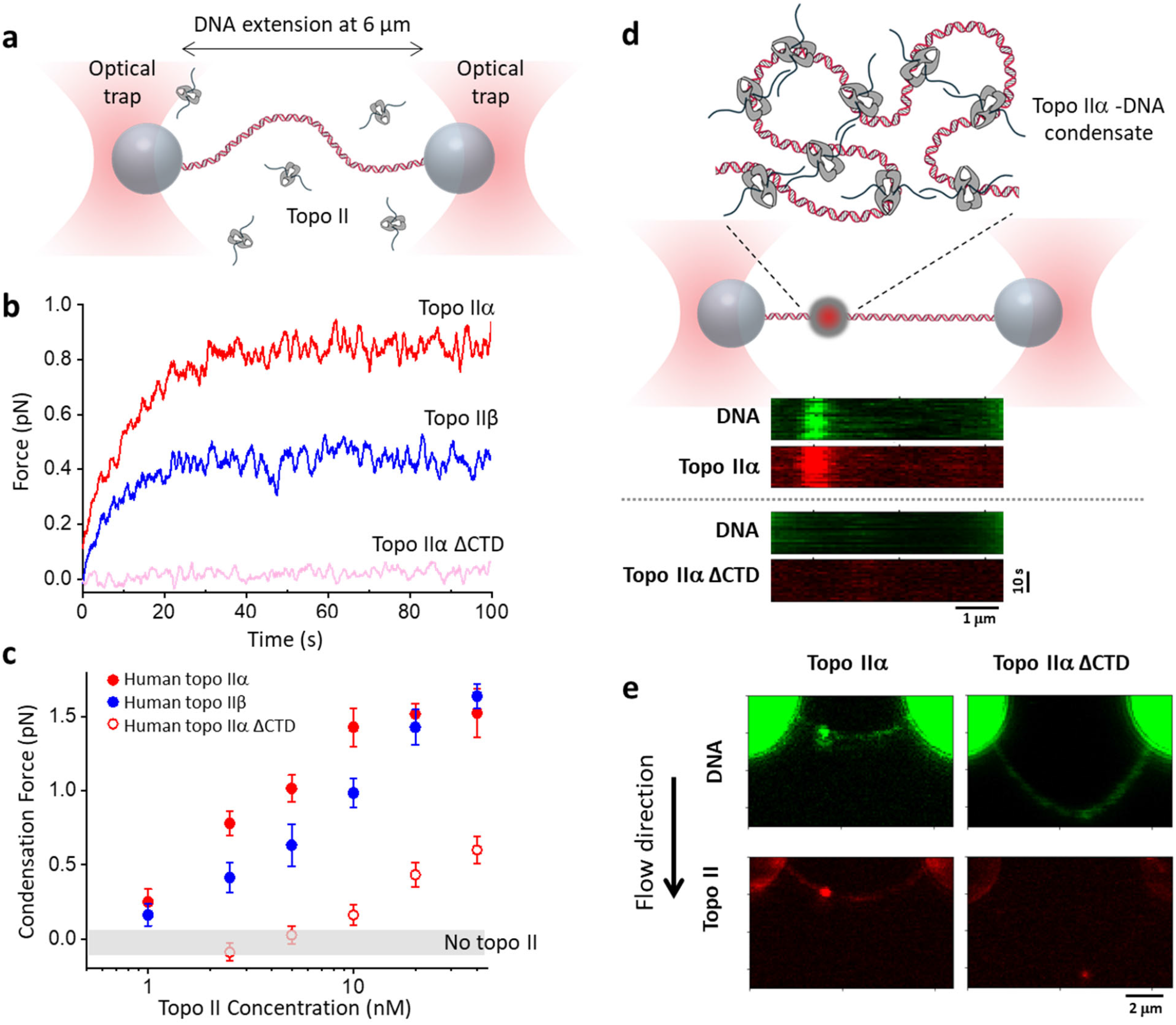
Topo IIα generates a driving force to condense DNA and form topo IIα-DNA condensates. **a**. Experimental scheme for monitoring the force generated during topo II condensation of DNA. A λ DNA molecule was tethered between two streptavidin-coated polystyrene beads held in two optical traps. The DNA extension was held at 6 μm, which resulted in a near-zero force applied to the DNA in the absence of topo II. **b**. Example traces for force generation during topo II condensation of DNA. The concentration of each topo II isoform was 2.5 nM. **c**. The final condensation force generated as a function of topo II concentration for three topo II isoforms. Each data point corresponds to an average of ∼20 individual traces, and the error bars indicate SDs. No topo control is also shown to provide the measurement uncertainty of the instrument. **d**. Topo IIα forms condensates on DNA. The top cartoon illustrates a topo IIα-DNA condensate formed along a λ DNA molecule held between two optical traps. The DNA extension was held at 8 μm, which resulted in a near-zero force applied to the DNA in the absence of topo II. Kymographs of confocal scans of both topo II and DNA were taken after allowing 5 nM AF488-labelled human topo IIα to condense DNA (visualized via Sytox Orange) for 70 s. **e**. Representative flow-stretch confocal images of AF488-human topo IIα and DNA stained by Sytox Orange. The flow stretches the DNA in the direction of the flow, which could reveal any large DNA loops associated with the condensate.

We found the condensation force to be highly topo II concentration-dependent, increasing with an increase in the topo II concentration (Fig. 5c). This concentration dependence reflects the critical concentration of the condensation phase transition being force-dependent (Supplementary Fig. 7). The condensation force reached a maximum of around 1.5 pN for both topo IIα and topo IIβ, although topo IIα can reach this maximum force at a lower concentration. Topo IIα ΔCTD condensation was ineffective at force generation, reaching only 0.5 pN under the highest concentration (40 nM) measured here, consistent with CTD-truncated topo IIα being ineffective at condensing DNA.

Collectively, our experiments demonstrate that both human topo IIα and topo IIβ can generate a maximum condensation force of about 1.5 pN, though topo IIα can reach this force at a lower topo II concentration. Although these forces are much lower than those generated by RNA polymerases during transcription^58-60^, they are somewhat larger than those previously measured for yeast condensin during loop extrusion (0.4 – 1.0 pN)^46^.

## Visualizing topo IIα-DNA condensates

If topo IIα condenses DNA after a phase transition, we expect this to lead to a phase separation along the DNA. Thus, DNA should exist in two phases: the DNA-only phase, and the condensate phase as a mixture of topo IIα and DNA. To test this prediction, we conducted an experiment similar to that used to measure the condensation force, except using AF488 labeled human topo IIα (5 nM) and Sytox Orange labelled DNA. Strikingly, we observed the formation of topo IIα-DNA condensates along the DNA (Fig. 5d). Frequently, we observed only a single condensate, with a strong intensity of the DNA signal overlapping with a strong topo IIα signal. In contrast, in the presence of 5 nM AF488-human topo IIα ΔCTD, we never detected any large condensates.

To exclude the possibility that the DNA signal resulted from a large DNA loop formed by a topo IIα cluster formed at the base of the loop, we flow-stretched the DNA, which could also extend any DNA in a loop in the direction of the flow, making such a DNA loop more visible (Fig. 5e). In the presence of 5 nM AF488-human topo IIα, we detected a shortened DNA path length between the trapped beads, consistent with DNA being sequestered in a condensate. No DNA loops were seen to emanate from the topo IIα cluster, further indicating that the body corresponded to a topo IIα-DNA condensate. In the presence of 5 nM human AF488-topo IIα ΔCTD, we always detected a much longer DNA path between the trapped beads and sometimes a very small condensate, indicating that the majority of DNA was not sequestered in any condensates (Fig. 5e).

## Discussion

In the present work, we uncovered an unsuspected new role for human topo IIα that functionally differentiates it from human topo IIβ. Human topo IIα is significantly more efficient than topo IIβ at condensing DNA or chromatin through a highly cooperative phase transition that is dependent on its unstructured CTD (Fig. 3). Importantly, condensation is driven by a process distinct from topo IIα’s catalytic activity and minimally interferes with its strand passage function (Fig. 4). Although we also found that human topo IIα is modestly more effective than human topo IIβ at decatenating chromatin, what truly distinguishes the enzyme from its counterpart is its effectiveness in condensing DNA.

Our findings suggest that topo IIα may directly help drive chromosome condensation during mitosis. This activity significantly expands the more established role of the enzyme in resolving supercoiling and crossovers that might result from loop extrusion by condensin I. We discovered that topo IIα’s DNA condensation force is slightly greater than the force yeast condensin generates during loop-extrusion (Fig. 5c), further supporting the notion that its condensation function is likely to be physiologically meaningful. Because each dimer of topo IIα contains two CTDs, each of which is thought to bind DNA and may also participate in inter-topo II interactions^17^, the enzyme may support multivalent contacts that assist with condensation (see Fig. 5d inset). DNA bending induced by topo IIα may further enhance these interactions by increasing the local topo IIα concentration. These inter-topo IIα interactions could bridge different regions of the same chromosome or connect sister chromatids, as suggested by studies of LLPS by topo II^17^. Significantly, our findings are consistent with a polymer-collapse phase transition that occurs at a much lower topo IIα concentration (100 pM or less), in contrast to the 100 nM topo IIα concentration required for LLPS^17^.

Previous studies have shown that topo IIα and condensin I localize to scaffold the axis of a mitotic chromosome, which runs along the length of a chromosome and appears to serve as a structural backbone that organizes chromatin loops and allows for the highly condensed linear arrangement of each sister chromatid^33,35,61,62^. Topo IIα molecules may localize to this axis because condensin I loops bring DNA segments into proximity, facilitating topo IIα-topo IIα interactions. Alternatively, inter-topo IIα interactions may help initiate the process, which is then supported by condensin I to help organize the disparate DNA segments that are being randomly brought into close local proximity by the topo IIα collective. Although topo IIα localizes to mitotic chromosomes in the nucleus, topo IIβ is found to be excluded from the nucleus and present mostly in the cytosol^16,62-64^. This indicates that topo IIβ concentration around DNA may be well below what is required to support condensation (the number of topo IIβ copies in cell is also estimated to be about 3.5 times lower than topo IIα copies during the G2-M phase^24^). Moreover, our measurements show that the critical concentration for efficient DNA condensation by topo IIβ is about 4 times higher than that for topo IIα (Fig. 3c). Combined, these factors give topo IIα more than 100-fold advantage in facilitating chromosome condensation compared to topo IIβ (SI Note). This finding may help explain why humans have two topo II isoforms. Similarly, because topo IIα is prone to condense DNA, it may need to be kept off the chromosome during non-replicative cell cycle phases such as G1 to permit chromatin to decondense and allow processes such as transcription to efficiently occur.

Our findings also have significant implications for understanding the consequences of chemotherapeutic drugs, such as etoposide and ICRF187, that target human topo IIs^48,65,66^. Existing drugs target the human topo IIα catalytic domain, which is shared between the two isoforms, leading to detrimental off-target side effects^67-69^. This suggests that there may be significant utility in identifying agents that specifically target the topo IIα CTD to induce mitotic failure. Although the unstructured nature of the topo IIα CTD makes drug design challenging, efforts are underway to identify specific antagonists of LLPS for other proteins with an intrinsically disordered domain^70,71^. What makes this more difficult is the lack of functional readout for a drug assay. Our assays for observing topo IIα-induced DNA condensation should be well-suited for testing drugs targeting the topo IIα CTD to disrupt its ability to condense DNA.

## ACKNOWLEDGEMENTS

We thank members of the Wang Laboratory for helpful discussion and comments. This work is supported by the National Institutes of Health grants R01GM136894 (to M.D.W.), R35CA263778 (to J.M.B.), T32GM008267 (to M.D.W.), and 5T32GM144272-03 (to J.M.B.). M.D.W. is a Howard Hughes Medical Institute investigator.

## AUTHOR CONTRIBUTIONS

M.W. and M.D.W. designed single-molecule assays. M.W. and M.V.W performed experiments. M.W., J.T.I., and M.D.W. analyzed data. C.B., J.H.L., J.J., and J.M.B. purified and characterized topoisomerases. M.W. prepared and characterized DNA constructs and chromatin substrates. R.M.F. purified histones. M.W. and M.D.W. wrote the initial draft, and all authors contributed to manuscript revisions. M.D.W. supervised the project.

## Code availability

Data analysis routines used to process and generate plots are available on Github: https://github.com/WangLabCornell/Wu_et_al.

## DATA AND MATERIAL AVAILABILITY

All data are available in the main text or the supplementary materials.

## References

1 Kouzine, F., Liu, J., Sanford, S., Chung, H.-J. & Levens, D. The dynamic response of upstream DNA to transcription-generated torsional stress. Nature Structural & Molecular Biology 11, 1092–1100, doi:10.1038/nsmb848 (2004).

2 Kouzine, F., Sanford, S., Elisha-Feil, Z. & Levens, D. The functional response of upstream DNA to dynamic supercoiling in vivo. Nature Structural & Molecular Biology 15, 146–154, doi:10.1038/nsmb.1372 (2008).

3 Naughton, C. et al. Transcription forms and remodels supercoiling domains unfolding large-scale chromatin structures. Nature Structural & Molecular Biology 20, 387–395, doi:10.1038/nsmb.2509 (2013).

4 Keszthelyi, A., Minchell, N. E. & Baxter, J. The causes and consequences of topological stress during DNA replication. Genes 7, 134 (2016).

5 Branzei, D. & Foiani, M. Maintaining genome stability at the replication fork. Nature Reviews Molecular Cell Biology 11, 208–219 (2010).

6 Ma, J., Bai, L. & Wang, M. D. Transcription Under Torsion. Science 340, 1580–1583, doi:10.1126/science.1235441 (2013).

7 Ma, J. et al. Transcription factor regulation of RNA polymerase’s torque generation capacity. Proceedings of the National Academy of Sciences 116, 2583–2588, doi:10.1073/pnas.1807031116 (2019).

8 Pommier, Y., Nussenzweig, A., Takeda, S. & Austin, C. Human topoisomerases and their roles in genome stability and organization. Nature Reviews Molecular Cell Biology 23, 407–427 (2022).

9 Nitiss, J. L. DNA topoisomerase II and its growing repertoire of biological functions. Nature Reviews Cancer 9, 327–337, doi:10.1038/nrc2608 (2009).

10 Pommier, Y., Sun, Y., Huang, S.-y. N. & Nitiss, J. L. Roles of eukaryotic topoisomerases in transcription, replication and genomic stability. Nature Reviews Molecular Cell Biology 17, 703–721, doi:10.1038/nrm.2016.111 (2016).

11 Lee, J. H. & Berger, J. M. Cell cycle-dependent control and roles of DNA topoisomerase II. Genes 10, 859 (2019).

12 Killian, J. L., Ma, J. & Wang, M. D. RNA polymerase as a torsional motor. (2021).

13 Berger, J. M., Gamblin, S. J., Harrison, S. C. & Wang, J. C. Structure and mechanism of DNA topoisomerase II. Nature 379, 225–232 (1996).

14 Champoux, J. J. DNA topoisomerases: structure, function, and mechanism. Annual review of biochemistry 70, 369–413 (2001).

15 Roca, J. & Wang, J. C. DNA transport by a type II DNA topoisomerase: evidence in favor of a two-gate mechanism. Cell 77, 609–616 (1994).

16 Linka, R. M. et al. C-terminal regions of topoisomerase II α and II β determine isoform-specific functioning of the enzymes in vivo. Nucleic Acids Research 35, 3810–3822 (2007).

17 Jeong, J., Lee, J. H., Carcamo, C. C., Parker, M. W. & Berger, J. M. DNA-Stimulated Liquid-Liquid phase separation by eukaryotic topoisomerase ii modulates catalytic function. eLife 11, e81786 (2022).

18 Gilroy, K. L. & Austin, C. A. The impact of the C-Terminal domain on the interaction of human DNA topoisomerase II α and β with DNA. PLOS ONE 6, e14693 (2011).

19 Akimitsu, N. et al. Enforced cytokinesis without complete nuclear division in embryonic cells depleting the activity of DNA topoisomerase IIα. Genes to Cells 8, 393–402 (2003).

20 McKinnon, P. J. Topoisomerases and the regulation of neural function. Nature Reviews Neuroscience 17, 673–679 (2016).

21 Lyu, Y. L. & Wang, J. C. Aberrant lamination in the cerebral cortex of mouse embryos lacking DNA topoisomerase IIβ. Proceedings of the National Academy of Sciences 100, 7123–7128, doi:10.1073/pnas.1232376100 (2003).

22 Akimitsu, N. et al. Induction of apoptosis by depletion of DNA topoisomerase IIα in mammalian cells. Biochemical and biophysical research communications 307, 301–307, doi:10.1016/S0006-291X(03)01169-0 (2003).

23 Heck, M. & Earnshaw, W. C. Topoisomerase II: A specific marker for cell proliferation. The Journal of cell biology 103, 2569–2581 (1986).

24 Woessner, R. D., Mattern, M. R., Mirabelli, C. K., Johnson, R. K. & Drake, F. H. Proliferation-and cell cycle-dependent differences in expression of the 170 kilodalton and 180 kilodalton forms of topoisomerase II in NIH-3T3 cells. Cell Growth Differ 2, 209–214 (1991).

25 Paulson, J. R., Hudson, D. F., Cisneros-Soberanis, F. & Earnshaw, W. C. in Seminars in cell & developmental biology. 7–29 (Elsevier).

26 Maeshima, K. & Eltsov, M. Packaging the genome: the structure of mitotic chromosomes. Journal of biochemistry 143, 145–153 (2008).

27 Kschonsak, M. & Haering, C. H. Shaping mitotic chromosomes: From classical concepts to molecular mechanisms. Bioessays 37, 755–766 (2015).

28 Samejima, K. et al. Mitotic chromosomes are compacted laterally by KIF4 and condensin and axially by topoisomerase IIα. Journal of Cell Biology 199, 755–770 (2012).

29 Baxter, J. & Aragón, L. A model for chromosome condensation based on the interplay between condensin and topoisomerase II. Trends in Genetics 28, 110–117, doi:10.1016/j.tig.2011.11.004 (2012).

30 Vos, S. M., Tretter, E. M., Schmidt, B. H. & Berger, J. M. All tangled up: how cells direct, manage and exploit topoisomerase function. Nature Reviews Molecular Cell Biology 12, 827–841 (2011).

31 Meczes, E. L., Gilroy, K. L., West, K. L. & Austin, C. A. The Impact of the Human DNA Topoisomerase II C-Terminal Domain on Activity. PLOS ONE 3, e1754, doi:10.1371/journal.pone.0001754 (2008).

32 Farr, C. J., Antoniou-Kourounioti, M., Mimmack, M. L., Volkov, A. & Porter, A. C. The α isoform of topoisomerase II is required for hypercompaction of mitotic chromosomes in human cells. Nucleic Acids Research 42, 4414–4426 (2014).

33 Shintomi, K. & Hirano, T. Guiding functions of the C-terminal domain of topoisomerase IIα advance mitotic chromosome assembly. Nature Communications 12, 2917 (2021).

34 Antoniou-Kourounioti, M., Mimmack, M. L., Porter, A. C. G. & Farr, C. J. The Impact of the C-Terminal Region on the Interaction of Topoisomerase II Alpha with Mitotic Chromatin. International Journal of Molecular Sciences 20, 1238 (2019).

35 Zhang, M. et al. Histone H2A phosphorylation recruits topoisomerase II α to centromeres to safeguard genomic stability. The EMBO Journal 39, e101863 (2020).

36 Lane, A. B., Giménez-Abián, J. F. & Clarke, D. J. A novel chromatin tether domain controls topoisomerase IIα dynamics and mitotic chromosome formation. Journal of Cell Biology 203, 471–486 (2013).

37 Sundararajan, S. et al. Methylated histones on mitotic chromosomes promote topoisomerase IIα function for high fidelity chromosome segregation. Iscience 26 (2023).

38 Austin, C. A. et al. Expression, Domain Structure, and Enzymatic Properties of an Active Recombinant Human DNA Topoisomerase IIβ (∗). Journal of Biological Chemistry 270, 15739–15746 (1995).

39 McClendon, A. K. et al. Bimodal recognition of DNA geometry by human topoisomerase IIα: preferential relaxation of positively supercoiled DNA requires elements in the C-terminal domain. Biochemistry 47, 13169–13178 (2008).

40 McClendon, A. K., Rodriguez, A. C. & Osheroff, N. Human Topoisomerase IIα Rapidly Relaxes Positively Supercoiled DNA: IMPLICATIONS FOR ENZYME ACTION AHEAD OF REPLICATION FORKS*. Journal of Biological Chemistry 280, 39337–39345, doi:10.1074/jbc.M503320200 (2005).

41 Seol, Y., Gentry, A. C., Osheroff, N. & Neuman, K. C. Chiral Discrimination and Writhe-dependent Relaxation Mechanism of Human Topoisomerase IIα*♦. Journal of Biological Chemistry 288, 13695–13703, doi:10.1074/jbc.M112.444745 (2013).

42 Le, T. T. et al. Synergistic coordination of chromatin torsional mechanics and topoisomerase activity. Cell 179, 619-631. e615 (2019).

43 Yurov, Y. B. Rate of DNA replication fork movement within a single mammalian cell. Journal of molecular biology 136, 339–342 (1980).

44 Conti, C. et al. Replication fork velocities at adjacent replication origins are coordinately modified during DNA replication in human cells. Molecular biology of the cell 18, 3059–3067 (2007).

45 Wang, W. et al. Genome-wide mapping of human DNA replication by optical replication mapping supports a stochastic model of eukaryotic replication. Molecular Cell 81, 2975-2988.e2976, doi:10.1016/j.molcel.2021.05.024 (2021).

46 Kim, E., Barth, R. & Dekker, C. Looping the genome with SMC complexes. Annual review of biochemistry 92, 15–41 (2023).

47 Chu, L. et al. The 3D topography of mitotic chromosomes. Molecular Cell 79, 902-916. e906 (2020).

48 Le, T. T. et al. Etoposide promotes DNA loop trapping and barrier formation by topoisomerase II. Nature chemical biology 19, 641–650 (2023).

49 Gao, X., Hong, Y., Ye, F., Inman, J. T. & Wang, M. D. Torsional stiffness of extended and plectonemic DNA. Physical review letters 127, 028101 (2021).

50 Daniels, B. C., Forth, S., Sheinin, M. Y., Wang, M. D. & Sethna, J. P. Discontinuities at the DNA supercoiling transition. Physical Review E—Statistical, Nonlinear, and Soft Matter Physics 80, 040901 (2009).

51 Forth, S. et al. Abrupt Buckling Transition Observed during the Plectoneme Formation of Individual DNA Molecules. Physical review letters 100, 148301, doi:10.1103/PhysRevLett.100.148301 (2008).

52 Hong, Y. et al. Optical torque calculations and measurements for DNA torsional studies. Biophysical Journal 123, 3080–3089, doi:10.1016/j.bpj.2024.07.005 (2024).

53 Mo, Y.-Y. & Beck, W. T. Association of Human DNA Topoisomerase IIα with Mitotic Chromosomes in Mammalian Cells Is Independent of Its Catalytic Activity. Experimental cell research 252, 50–62, doi:10.1006/excr.1999.4616 (1999).

54 Sheinin, M. Y., Li, M., Soltani, M., Luger, K. & Wang, M. D. Torque modulates nucleosome stability and facilitates H2A/H2B dimer loss. Nature Communications 4, 2579 (2013).

55 Lee, J. et al. Chromatinization modulates topoisomerase II processivity. Nature Communications 14, 6844, doi:10.1038/s41467-023-42600-z (2023).

56 Rouches, M. N. & Machta, B. B. Polymer Collapse & Liquid-Liquid Phase-Separation are Coupled in a Generalized Prewetting Transition. ArXiv (2024).

57 Strick, R., Strissel, P. L., Gavrilov, K. & Levi-Setti, R. Cation–chromatin binding as shown by ion microscopy is essential for the structural integrity of chromosomes. The Journal of cell biology 155, 899–910 (2001).

58 Yin, H. et al. Transcription Against an Applied Force. Science 270, 1653–1657, doi:10.1126/science.270.5242.1653 (1995).

59 Wang, M. D. et al. Force and Velocity Measured for Single Molecules of RNA Polymerase. Science 282, 902–907, doi:10.1126/science.282.5390.902 (1998).

60 Hodges, C., Bintu, L., Lubkowska, L., Kashlev, M. & Bustamante, C. Nucleosomal Fluctuations Govern the Transcription Dynamics of RNA Polymerase II. Science 325, 626–628, doi:10.1126/science.1172926 (2009).

61 Nielsen, C. F., Zhang, T., Barisic, M., Kalitsis, P. & Hudson, D. F. Topoisomerase IIα is essential for maintenance of mitotic chromosome structure. Proceedings of the National Academy of Sciences 117, 12131–12142 (2020).

62 Meyer, K. N. et al. Cell cycle–coupled relocation of types I and II topoisomerases and modulation of catalytic enzyme activities. The Journal of cell biology 136, 775–788 (1997).

63 Grue, P. et al. Essential mitotic functions of DNA topoisomerase IIα are not adopted by topoisomerase IIβ in human H69 cells. Journal of Biological Chemistry 273, 33660–33666 (1998).

64 Christensen, M. O. et al. Dynamics of human DNA topoisomerases IIα and IIβ in living cells. The Journal of cell biology 157, 31–44 (2002).

65 Baldwin, E. & Osheroff, N. Etoposide, topoisomerase II and cancer. Current Medicinal Chemistry-Anti-Cancer Agents 5, 363–372 (2005).

66 Classen, S., Olland, S. & Berger, J. M. Structure of the topoisomerase II ATPase region and its mechanism of inhibition by the chemotherapeutic agent ICRF-187. Proceedings of the National Academy of Sciences 100, 10629–10634 (2003).

67 Nitiss, J. L. Targeting DNA topoisomerase II in cancer chemotherapy. Nature Reviews Cancer 9, 338–350, doi:10.1038/nrc2607 (2009).

68 Pommier, Y., Leo, E., Zhang, H. & Marchand, C. DNA Topoisomerases and Their Poisoning by Anticancer and Antibacterial Drugs. Chemistry & Biology 17, 421–433, doi:10.1016/j.chembiol.2010.04.012 (2010).

69 Azarova, A. M. et al. Roles of DNA topoisomerase II isozymes in chemotherapy and secondary malignancies. Proceedings of the National Academy of Sciences 104, 11014–11019, doi:10.1073/pnas.0704002104 (2007).

70 Conti, B. A. & Oppikofer, M. Biomolecular condensates: new opportunities for drug discovery and RNA therapeutics. Trends in Pharmacological Sciences 43, 820–837, doi:10.1016/j.tips.2022.07.001 (2022).

71 Zhang, Y., Jin, C., Xu, X., Guo, J. & Wang, L. The role of liquid-liquid phase separation in the disease pathogenesis and drug development. Biomedicine & Pharmacotherapy 180, 117448, doi:10.1016/j.biopha.2024.117448 (2024).

