## Supplementary Materials for "Human Topoisomerase IIα Promotes Chromatin Condensation Via a Phase Transition"

### Method Details

#### Protein Expression and Purification

All topoisomerase II (topo II) proteins (Supplementary Fig.1), including full-length and C-terminal domain (CTD)-truncated versions of human topo IIα, topo IIβ, and yeast topo II, were expressed from *S. cerevisiae* strain BCY123, as described previously[1-5]. Cells were lysed by cryogenic grinding, and the resulting lysate was clarified by centrifugation and filtering through 1.1 um syringe filters (Thermo Scientific #722-2000). Clarified lysate was loaded onto a 5 ml HisTrap HP (GE Healthcare) nickel-chelating Sepharose column and then a HiTrap SP (GE Healthcare) cation exchange column for further purification. The protein was then incubated with His6-tagged TEV protease (QB3 MacroLab) to remove the His tag, loaded onto a 1 mL HisTrap HP column to remove the protease and further purified on a gel-filtration column (S-400, GE Healthcare). Finally, the purified protein was concentrated, spiked with glycerol to 30%, and flash frozen for storage at -80 °C.

Human histone octamers were purified from HeLa-S3 cells[2,4]. Nuclei were extracted from HeLa-S3 cell pellets (National Cell Culture Center), and core histones were purified using a hydroxyapatite Bio-Gel HTP gel (Bio-Rad Laboratories # 130-0420) slurry. The purified core histones were concentrated and flash frozen for storage at -80 °C.

#### Fluorescence Labeling of Topo II

Purified topo II proteins were labeled with Alexa Fluor 488 (AF488) through NHS-Ester labeling[5]. In brief, these purified topo II proteins were incubated with AF488 NHS ester molecules with a molar ratio of 1:1.5 for dye labeling. The reaction mixture was then loaded onto a Zeba Spin Desalting Column (7 kDa MWCO, Thermo Scientific #89890) two consecutive times to remove the free dye. The purified AF488 labeled proteins were then concentrated and flash-frozen for storage at -80 °C.

#### DNA Template Construction

The DNA template used in the topo II induced DNA compaction experiments was torsionally constrained, containing a 12,688-bp DNA center segment and a ~500-bp multi-labeled adapter at each end[3,4]. The 12,688-bp DNA center segment was PCR amplified from λ-DNA (NEB, N3011S), then double digested with AvaI (NEB, R0152S) and BssSI-v2 (NEB, R0680S) to produce the unique overhangs for ligation. To make the 500-bp multi-biotin-labeled and multi-digoxigenin-labeled adapters, we performed PCR amplification from plasmid pNFRTC (pMDW111) with either 24% of dATP replaced by biotin-14-dATP or 24% of dTTP replaced by digoxigenin-11-dUTP, followed by restriction enzyme digestion with BssSI and AvaI, respectively. The ~500-bp multi-labeled adapters with unique overhangs were then ligated to the 12,688 bp center segment. The ligation product, 12.7 kb lambda template, was gel purified and aliquots were stored at -20 °C.

For nucleosome assembly, we used a torsionally constrained 12.7 kb 64-mer DNA template composed of 64 tandem repeats of a 197-bp sequence containing a 601 nucleosome positioning element (NPE), flanked by a ~500-bp multi-labeled adapters at each end[2,4]. The 64-mer center segment was created by double digesting the plasmid p197NRL-64ex (pMDW108) with BstXI (NEB, R0113S) and BglI (NEB, R0143S), followed by gel purification. The 500-bp multi-labeled adapters were prepared from pNFRTC as described above using a different primer sequence and digested with either BstXI for a multi-biotin-labeled adapter or BglI for multi-digoxigenin-labeled adapter. The multi-labeled adapters with unique overhangs were ligated to the center segment, to produce 12.7 kb 64-mer template used for nucleosome assembly.

**Nucleosome Assembly**

Nucleosome arrays were assembled by salt dialysis using the 12.7 kb 64-mer DNA construct and purified HeLa histone octamers[2,6,7]. The 12.7 kb 64-mer template and histone octamers were mixed at different molecular ratios of histone octamers to 601 NPE (1.3:1 to 1.5:1), followed by a gradient NaCl salt dialysis from 2 M to 0 M over 24 hours at 4°C. To avoid nucleosome over-assembly, an equal mass of 147-bp random sequence competitor DNA was included in the assembly reactions. The quality and saturation level of the assembled nucleosome arrays were initially assessed by gel electrophoresis using a 0.7% native agarose gel, and then further rigorously characterized by nucleosome stretching using optical tweezers and twisting using magnetic tweezers[4].

**Single-Molecule Sample Chamber Preparation**

Single-molecule sample chambers were formed by assembling two nitrocellulose-coated microscope coverslips using inert silicone high vacuum grease4. The coverslips were prepared by coating a thin film of nitrocellulose solution (1-2% collodion in Amyl acetate) onto a pre-cleaned coverslip using a spin coater. This was followed by 95% ethanol incubation, baking at 80°C for further hardening, water rinses, and drying.

Experiments described in Figs. 1-4 were performed with DNA tethers that were torsionally constrained between a streptavidin-coated magnetic bead via multiple biotin-streptavidin interactions and the surface of the sample chamber via multiple digoxygenin-anti-digoxygenin interactions. To form single DNA tethers in a sample chamber, the surface of the chamber was functionalized with anti-digoxygenin (Vector Labs MB-7000), passivated with β-Casein (Sigma C6905), incubated with 2 pM 12.7 kb lambda template or 7 pM 12.7 kb 64-mer nucleosome array, followed by incubation with paramagnetic beads (Invitrogen 65601). To form double chromatin tethers in the sample chamber, the surface of the chamber was first functionalized and passivated as described above. A pre-incubation of the nucleosome arrays and streptavidin coated magnetic beads was performed and then introduced into the sample chamber for further incubation. This was followed by an incubation with the nicking enzyme, Nt.BsmAl (NEB, R0121S) to prevent torsional buildup within each DNA molecule of a double chromatin tether. To enable the comparison of topo II activity between single and double chromatin fibers in Fig.1, both single chromatin chambers and double chromatin chambers were prepared in the same way with only difference being a lack of nicking enzyme in the single chromatin chambers. The pre-incubation step for all chambers was performed for 15 minutes at an optimized molecular ratio of nucleosome array and magnetic bead. This resulted in a similar amount of single and double chromatin tethers present in the respective chambers. Immediately prior to the surface passivation step with β-Casein, magnetic beads coated with 500 bp multiple biotin and digoxigenin labeled DNA were bound to the surface of the sample chamber to serve as fiducial markers.

Prior to an experiment, a buffer exchange was done and, unless stated otherwise, all experiments were carried out in the topo reaction buffer (10 mM Tris-HCl pH 8.0, 50 mM NaCl, 50 mM KCl, 3 mM $MgCl_2$, 0.1 mM EDTA, 1 mM DTT, 0.5 mM TCEP, 1 mM ATP, and 1.5 mg/mL β-Casein) in a soundproof room at a temperature of 23°C.

**Topo II Activity and Condensation Assay on Magnetic Tweezers (MT)**

MT experiments described in Figs. 1-4 for assessing catalytic activity and DNA condensation of topo II were carried on a home-built MT setup, allowing real-time supercoiling and extension monitoring under a constant force of 0.5 pN for multiple molecules simultaneously[2-4].

Before introducing topo II to the sample chamber, the chromatin fiber substrates were assayed to determine the nucleosome composition, geometry, and stability. The substrates were twisted by rotation of the magnet to obtain the initial extension versus turns relation for each tether (Supplementary Fig. 3). These initial extension versus turn relations of single chromatin substrates were fit to five-piece functions to obtain peak extension and "buckling-like" transitions for both negative and right turns, allowing characterization of the quality and saturation level of the nucleosome arrays. We selected traces with good nucleosome composition and an extension consistent with 50 ± 5 nucleosomes[2,4]. For the double chromatin fibers, we selected traces with small anchor separations at both ends and an extension consistent with 50 ± 5 nucleosomes[2].

Topo II catalytic activity measurements in Fig.1 and Fig.4 were performed with a continuous winding assay[2]. After introducing topo II, the chromatin substrates were supercoiled by continuously rotating the magnets at 3.6 turns/s for 1000 turns. While the substrates were continuously twisted, the extensions were monitored to assay the ability of topo II to remove the added supercoils.

DNA condensation measurements of Topo II in Figs. 2 and 4 were also performed on MT. After introducing topo II, the chromatin substrate extension was monitored to track the compaction of the tether. In Fig. 2, the magnets were quickly rotated at 10 turn/s for 50 (Fig. 2b) or 70 (Fig. 2d) turns to introduce supercoiling into the tethers. The inability to significantly decrease the tether extension and the subsequent recovery of the extension decrease indicates topo II catalytic activity.

**Minimum Extension of Naked DNA and Chromatin Tethers on MT**

DNA and chromatin tethers were formed on MT by surface immobilization to the sample chamber at one end and attaching to a 1 μm streptavidin coated magnetic beads at the other end. Since the magnetic beads were coated by streptavidin, the DNA or chromatin tethers were randomly attached on the surface of the beads, which can result in a non-zero extension of the tethers when the beads have touched the surface. This extension values ranges from 0 to 0.5 μm (the radius of the magnetic beads). We measured the average extension of tethers when the DNA or chromatin tethers were wound down to the surface by adding +150 turns. The measured average extension represented the minimum attainable value for the instrument, which was ~0.13 μm for both DNA and chromatin tethers. The minimum extension was further normalized by the extension of naked DNA and chromatin substrate under the experimental force 0.5pN, shown as gray dash line in Fig.3a and 3b.

**Estimation of Turns Relaxed in a Continuous Winding Experiment**

To estimate the number of turns relaxed in a continuous winding assay in Fig.4, we considered the final extension difference between 0 turn added (brown) vs 2160 turns added at a rate of 3.6 turns/s (red). The final extension difference for single substrates in Fig.4a is ~0.2 μm, which corresponding to ~50 turns based on the extension vs turns relation of a single chromatin substrate (Supplementary Fig.3a). We estimated that there is ~50 turns out of a total of 2160 turns added not being relaxed, therefore topo IIα relaxed 2110 turns, which is 98% turns added during continuous winding assay. Similarly, the final extension difference between 0 turn added (brown) and 2160 turns added (red) for double substrates in Fig.4b is ~0.1 μm, which corresponding to ~20 turns based on the extension vs turns relation of a double chromatin substrate. We estimated that there is ~20 turns out of a total of 2160 turns added not being relaxed, therefore topo IIα relaxed 2140 turns, which is 99% turns added during continuous winding assay.

**Force Generation and Topo II-DNA Bio-condensates visualization using Lumicks C-trap**

Force generation by topo II-DNA condensation (Fig. 5) was measured using LUMICKS C-trap. In a microfluidic flow cell, two 4.3-µm diameter streptavidin-coated polystyrene beads (Spherotech) were trapped in the bead channel and were subsequently moved to the DNA channel containing biotinylated λ-DNA (LUMICKS) for tether formation. The DNA tether was moved to a third channel with topo reaction buffer and was stretched to ensure that the tether only contained a single DNA molecule based on its force-extension curve. The tether, held at a constant extension of 6 µm, was then moved to the protein channel containing unlabeled topo II at various concentrations from 1 nM-40 nM in topo reaction buffer to monitor the force generation during the formation of topo II-DNA condensates.

To visualize the formation of topo II-DNA bio-condensates, we did similar experiments as described above, except using 5 nM AF488 labeled topo IIs and 10 nM Sytox Orange to visualize DNA. The tether, held at a constant extension of 8 µm, was then moved into the protein channel. AF488 labeled topo IIs were excited with a 488 nm laser and the DNA labeled with Sytox Orange was excited with a 532 nm laser. Kymographs of topo and Sytox labeled DNA were recorded at 2 lines/s during the formation of the bio-condensates. After the formation of the topo II-DNA bio-condensates, we flow stretched the tether by applying a flow rate of ~ 1 mm/s perpendicular to the tether and sequentially imaged AF488 labeled topo IIs and Sytox labeled DNA.

**Compaction Rate During Topo II Compaction of DNA**

The compaction rate was obtained by fitting individual extension versus time curves with a single exponential decay function (Supplementary Fig.5):

$$y = (1 - c) * e^{-bt} + c,$$

which yielded *b* as the compaction rate and *c* as the final compacted extension.

#### Condensation Efficiency for Topo II Condensation of DNA

The condensation efficiency $(1/[\text{topo II}]_c)$ , as shown in Fig. 3c, was calculated from the critical concentration for phase transition as topo II condensed DNA. We saw a sharp reduction in the final extension when topo II concentration reached a critical value, indicating cooperative binding on DNA and indicative of a phase transition. The Hill equation is used to fit the plots in Figs. 3a and 3b is:

$$y = 1 + (a - 1) * \frac{[\text{topo}]^n}{{[\text{topo}]_c}^n + [\text{topo}]^n},$$

where *a* is the normalized extension at maximum compaction, $[\text{topo}]_c$ is the critical topo II concentration at which the extension is reduced by half, and *n* is the Hill coefficient.

**Estimation of Human Topo IIα and Topo IIβ Association with Chromosome During Mitosis**

To estimate the concentrations of human topo IIα and topo IIβ during mitosis, we consider the following published observations. Human cells express roughly a million, $10^6$ copies of topo IIα and topo IIβ[8,9]. While topo IIα expression is cell-cycle regulated, with ~3.5 fold higher in G2 and M phases than in G1 and S phases, topo IIβ is present at uniform levels throughout the cell[10]. Moreover, topo IIα and topo IIβ show very different localization patterns in mitosis, with topo IIα localizing to the mitotic chromosomes in the nucleus, topo IIβ excluded from the nucleus and present mostly in the cytosol[11-14]. Since the human cell nucleus volume is roughly 10% of the human cell volume, the topo IIα concentration is estimated to be at least 35-fold higher than topo IIβ in the nucleus during mitosis.

Moreover, our measurements show that the critical concentration for topo IIβ phase transition is about 4 times higher than that for topo IIα (Fig. 3c). Thus, topo IIα has more than a 100-fold advantage over topo IIβ in facilitating chromosome condensation during mitosis.

### SI Figures

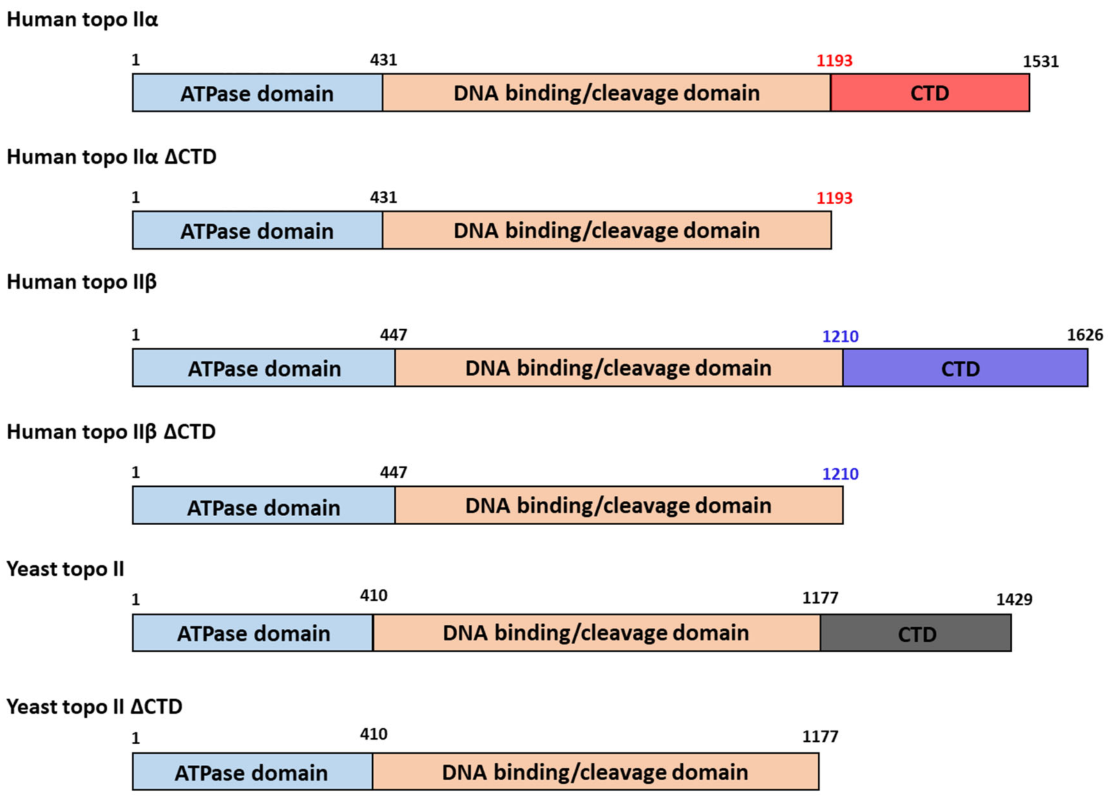


**Supplementary Fig. 1. Schematic synopsis of topoisomerase II proteins studied.**

Domain organization of the topoisomerase II (topo II) proteins used in this study. The eukaryotic topo II proteins from human and yeast share a conserved ATPase domain and DNA binding/cleavage domains and differ in the C-terminal domain (CTD). The topo II ΔCTD mutant proteins have the CTD domain completely truncated.

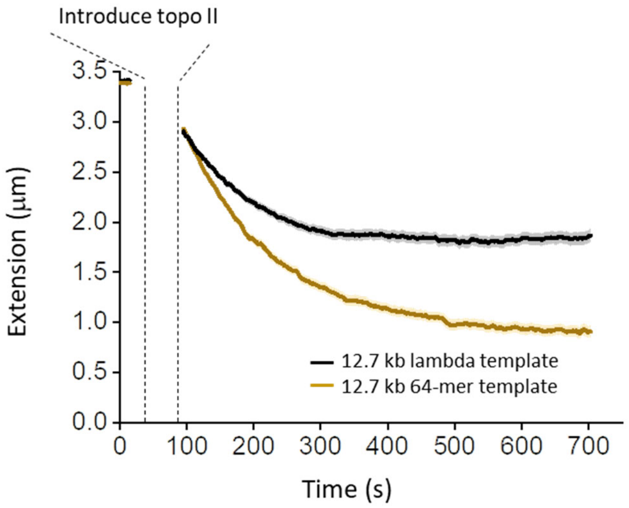


**Supplementary Fig. 2. Sequence dependence of topo II-induced DNA compaction.**

DNA extension *versus* time in the presence of 80 pM yeast topo II on 12.7 kb lambda template with a random non-repetitive sequence and 12.7 kb 64mer template with 64 repeats of the nucleosome positioning element. We observe some difference between the compaction of two DNA substrates that have the same DNA length but different sequences, suggesting that there is some sequence dependence in topo II-induced DNA compaction. Each curve represents the average of ~56 individual traces, with the shaded region indicating the SEM.

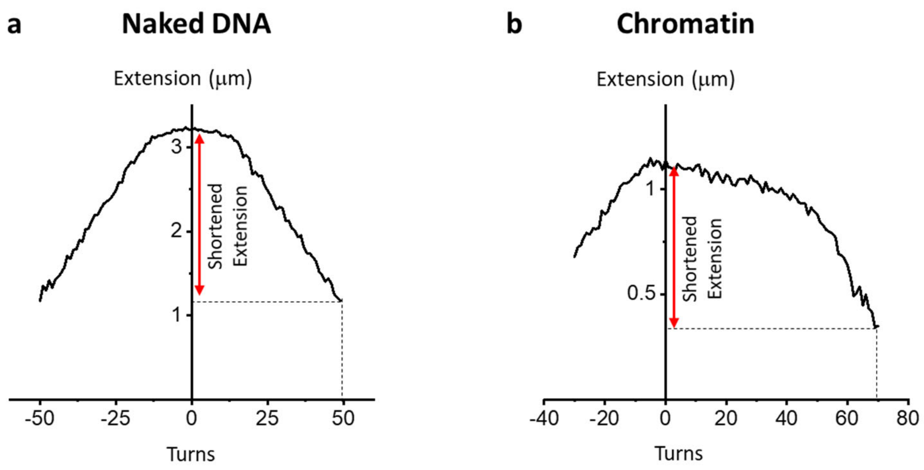


**Supplementary Fig. 3. Extension *versus* turns relation of DNA and Chromatin.**

**a.** Representative trace of the extension *versus* turns relation of naked DNA under 0.5 pN. In the absence of topo II, naked DNA is supercoiled by rotating the MT magnets. At (+) 50 turns, the extension of the DNA is significantly shortened due to the formation of plectonemes.

**b.** Representative trace of the extension *versus* turns relation of a chromatin substrate under 0.5 pN. In the absence of topo II, the chromatin substrate is supercoiled by rotating the MT magnets. At (+) 70 turns, the extension of the chromatin substrate is significantly shortened as the chromatin substrate also undergoes a buckling-like transition.

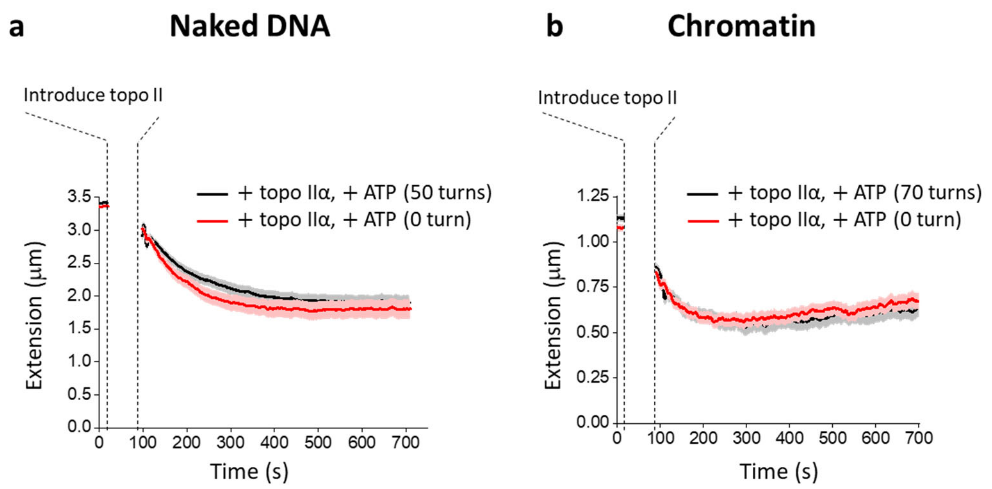


**Supplementary Fig. 4. An initial brief activity check does not interfere with topo II-induced DNA compaction.**

**a.** DNA extension *versus* time in the presence of human topo IIα with (50 turns added, black) and without (0 turn added, red) an initial activity check. We observe no difference in DNA compaction, indicating a brief activity check does not interfere with the compaction process. Experiments were conducted under a condition where the final extension was at about 50% of the initial extension. Each curve represents the average of ~60 individual traces, with the shaded region indicating the SEM.

**b.** Same as (a) except with the chromatin substrates by adding 70 turns for an initial activity check. Each curve represents the average of ~66 individual traces, with the shaded region indicating the SEM.

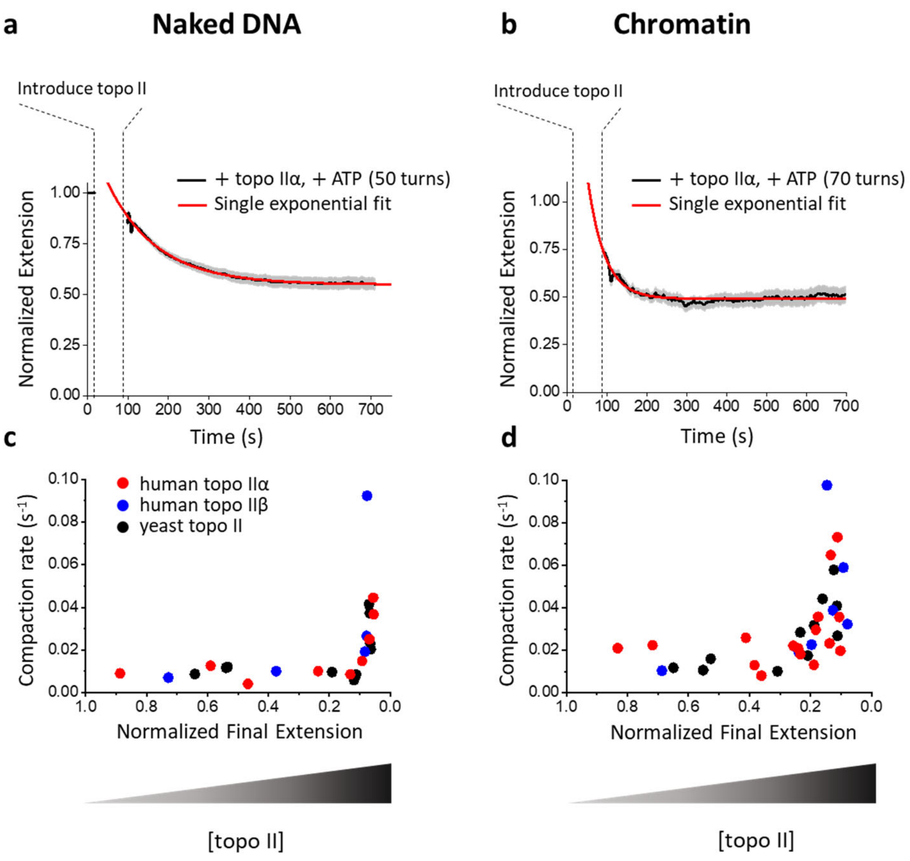


**Supplementary Fig. 5. Single exponential fit and compaction rate of topo II-induced compaction.**

**a**. Representative single exponential fit of normalized extension *versus* time of human topo IIα induced compaction from Fig. 2b (Methods): $y = (1 - c) * e^{-bt} + c$, yielding *b* as the compaction rate and *c* as the final compacted extension.

**b**. Same as (a) except with a chromatin substrate from Fig. 2d.

**c**. Compaction rate versus normalized final compacted DNA extension from the topo IIs: human topo IIα (60-200 pM), topo IIβ (400 pM-5 nM), yeast topo II (60-200 pM). For all three topo II isoforms, the final compacted DNA extension decreased with an increase in topo II concentration, indicating the extent of the compaction increasing with topo II concentration. However, the compaction rate remained constant at ~0.01 $s^{-1}$ before reaching the maximum compaction with the smallest final extension. After reaching the maximum compaction, the compaction rate increased with further topo II concentration. Each data point corresponds to a fitted value for the average of ~70 individual traces.

**d**. Same as (c) except with a chromatin substrate: human topo IIα (60-200 pM), topo IIβ (200 pM-1 nM), yeast topo II (60-250 pM). Each data point corresponds to a fitted value for the average of ~30 individual traces.

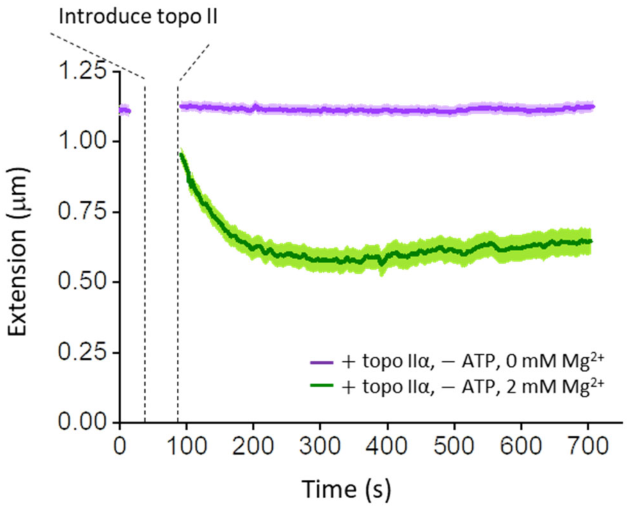


**Supplementary Fig. 6. Topo II-induced DNA compaction requires $Mg^{2+}$.**

Chromatin extension *versus* time in the presence of human topo IIα with (green) and without (purple) $Mg^{2+}$. Experiments were conducted under a condition where the final extension was at about 50% of the initial extension (in the presence of 0 or 2 mM $Mg^{2+}$). Each curve represents the average of ~62 individual traces, with the shaded region indicating the SEM.

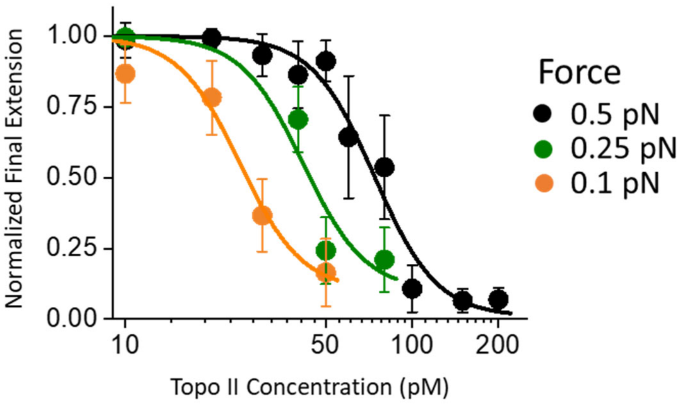


**Supplementary Fig. 7. Force dependence of topo II-induced DNA compaction.**

Normalized final DNA extension as a function of yeast topo II concentration under various forces, 0.5 pN, 0.25 pN and 0.1 pN. Topo II-induced DNA compaction is a force-dependent process: the smaller the force that was applied to the DNA, the more readily topo II can compact DNA and the lower the critical concentration required to induce a phase transition. Each normalized final extension data point represents an average of ~80 individual traces, with error bars indicating SDs.

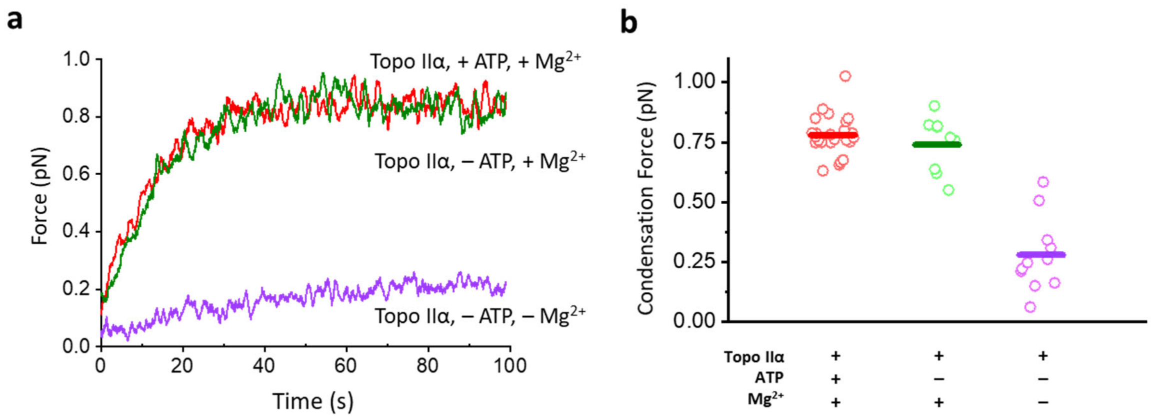


**Supplementary Fig. 8. Force generation by topo II on single DNA requires $Mg^{2+}$ but not ATP.**

**a.** Representative traces of force generation during human topo IIα condensation of DNA with (red) and without (green) ATP, as well as without ATP and $Mg^{2+}$ (purple). The concentration of topo IIα was 2.5 nM.

**b.** The final condensation force generated by topo IIα. The horizontal lines represent the mean force, while individual measurements from ~ 15 traces are shown as open circles.

#### References


1. Lee, J.H., Wendorff, T.J. & Berger, J.M. Resveratrol: A novel type of topoisomerase II inhibitor. *Journal of Biological Chemistry* **292**, 21011-21022 (2017).
2. Le, T.T. et al. Synergistic coordination of chromatin torsional mechanics and topoisomerase activity. *Cell* **179**, 619-631 e15 (2019).
3. Le, T.T. et al. Etoposide promotes DNA loop trapping and barrier formation by topoisomerase II. *Nat Chem Biol* **19**, 641-650 (2023).
4. Lee, J. et al. Chromatinization modulates topoisomerase II processivity. *Nat Commun* **14**, 6844 (2023).
5. Jeong, J., Lee, J.H., Carcamo, C.C., Parker, M.W. & Berger, J.M. DNA-Stimulated Liquid-Liquid phase separation by eukaryotic topoisomerase ii modulates catalytic function. *eLife* **11**, e81786 (2022).
6. Brower-Toland, B. et al. Specific contributions of histone tails and their acetylation to the mechanical stability of nucleosomes. *J Mol Biol* **346**, 135-46 (2005).
7. Huynh, V.A., Robinson, P.J. & Rhodes, D. A method for the in vitro reconstitution of a defined "30 nm" chromatin fibre containing stoichiometric amounts of the linker histone. *J Mol Biol* **345**, 957-68 (2005).
8. Heck, M. & Earnshaw, W.C. Topoisomerase II: A specific marker for cell proliferation. *The Journal of cell biology* **103**, 2569-2581 (1986).
9. Padget, K., Pearson, A. & Austin, C. Quantitation of DNA topoisomerase IIα and β in human leukaemia cells by immunoblotting. *Leukemia* **14**, 1997-2005 (2000).
10. Woessner, R.D., Mattern, M.R., Mirabelli, C.K., Johnson, R.K. & Drake, F.H. Proliferation-and cell cycle-dependent differences in expression of the 170 kilodalton and 180 kilodalton forms of topoisomerase II in NIH-3T3 cells. *Cell Growth Differ* **2**, 209-214 (1991).
11. Grue, P. et al. Essential mitotic functions of DNA topoisomerase IIα are not adopted by topoisomerase IIβ in human H69 cells. *Journal of Biological Chemistry* **273**, 33660-33666 (1998).
12. Linka, R.M. et al. C-terminal regions of topoisomerase II α and II β determine isoform-specific functioning of the enzymes in vivo. *Nucleic Acids Research* **35**, 3810-3822 (2007).
13. Christensen, M.O. et al. Dynamics of human DNA topoisomerases IIα and IIβ in living cells. *The Journal of cell biology* **157**, 31-44 (2002).
14. Meyer, K.N. et al. Cell cycle–coupled relocation of types I and II topoisomerases and modulation of catalytic enzyme activities. *The Journal of cell biology* **136**, 775-788 (1997).
